# A standardized naturalistic audio stimuli database with unsupervised labeling

**DOI:** 10.64898/2026.04.16.718910

**Authors:** Anas Al-Naji, Ricarda I. Schubotz, Anoushiravan Zahedi

**Author notes:** **Corresponding Author:** Correspondence concerning this article should be addressed to Anas Al-Naji & Anoushiravan Zahedi.

## Abstract

Research in cognitive neuroscience has relied on simple, highly controlled stimuli due to the difficulty in developing standardized, ecologically valid stimulus sets. However, there is a consensus that using ecologically valid stimuli is imperative to generalize results beyond controlled laboratory settings. The current study introduces a naturalistic audio stimulus database, consisting of short, recognizable, and emotionally rated stimuli. To create such a database, the current study collected 291 audio files from a wide range of sources. 361 participants rated the audio clips on emotionality, arousal, and recognizability, and subsequently freely described the audios by typing what they believed the sound to be. The text responses of the participants were embedded and clustered using an unsupervised machine-learning algorithm to derive a participant-grounded organization of auditory object categories. The results indicate audio clips were easily recognizable, while emotionality and arousal ratings showed broad variability, making the database suitable for diverse experimental needs. Furthermore, the final database comprises 10 distinct semantic categories, providing a diverse set of auditory stimuli.

## 1 Introduction

Research in cognitive neuroscience has traditionally relied on simple, highly standardized, and often abstract stimuli. The rationale for using such stimuli is to minimize experimental variability by controlling for potential confounds that more complex stimuli might introduce (Rust et al., 2005). However, the human brain evolved to process ecological sensory environments, which are rich and dynamic in nature (Sonkusare et al., 2019). Although an increasing number of studies using visual stimuli are incorporating more naturalistic materials into their designs (e.g., Bartels & Zeki, 2003; Hasson et al., 2004; Kauttonen et al., 2018; Lahnakoski et al., 2012), the dearth of ecologically valid auditory stimulus sets has resulted in continuing the use of simplified auditory stimuli. This gap in the field is mainly due to the challenges unique to the auditory domain; these challenges include, but are not restricted to, the absence of a universally agreed-upon definition of auditory objects and categories (Bizley & Cohen, 2013), the inherently temporal nature of sound (Griffiths & Warren, 2004), and the difficulty of recognizing and categorizing natural auditory sources (Bizley & Cohen, 2013; Zhai et al., 2020). Hence, although the necessity has been felt in the field (Stevenson & James, 2008), it is challenging to develop standardized, ecologically valid auditory stimulus databases suitable for the precise demands of cognitive neuroscience.

The definition of auditory objects remains debated in research. Early definitions described auditory objects as coherent constructs that emerge when the brain integrates acoustic features across time and frequency (i.e., timbre, pitch, and loudness), analogous to vision, where edges and contours can be combined to form visual objects (Griffiths & Warren, 2004). The current consensus, however, is that auditory objects represent the perceptual units over which the auditory system organizes the acoustic world (Bizley & Cohen, 2013). Bizley and Cohen (2013) emphasize the hierarchical nature of auditory object definition, where low-level features combine into objects, which themselves can be organized into increasingly abstract categories, and the categories can, in turn, be nested inside more general categories. For example, a dog’s bark or a bird’s call falls under broader categories such as mammals or birds, which together belong to the general category of living things. Because auditory objects and categories function as the fundamental building blocks of perception, carefully standardized data-driven databases that capture these distinctions are essential for advancing auditory cognition research.

Besides being difficult to standardize, auditory stimuli typically take longer to recognize than visual stimuli (Zhai et al., 2020), meaning that audio stimulus databases often include relatively long clips (e.g., The International Affective Digitized Sounds, IADS for short: 6 seconds). This poses a challenge for neuroimaging research, as longer stimulus durations reduce the number of trials that can be presented, thus lowering the overall signal-to-noise ratio (SNR; Boudewyn et al., 2017; Gonzalez-Moreno et al., 2014; Thigpen et al., 2016). Given the critical importance of SNR in cognitive neuroscience, there is a clear need for databases containing short yet still recognizable audio stimuli.

Although numerous auditory datasets exist (Fonseca et al., 2022; Gemmeke et al., 2017; Piczak, 2015; Salamon et al., 2014), most originate from computer science research and are primarily designed for developing and testing machine learning algorithms rather than for behavioral or biological studies. While such datasets are valuable for providing semantically variable stimuli for studying hierarchical auditory processing in the brain (de Heer et al., 2017), to the best of our knowledge, they mostly lack standardization along psychological dimensions, such as emotionality. This is in contrast to visual stimulus databases (Bradley & Lang, 2007; Brodeur et al., 2010; Marchewka et al., 2013), where stimuli are usually normed along dimensions such as emotionality, recognizability, and naming agreement. These characteristics are particularly valuable, as stimuli that evoke stronger emotions, for example, may elicit different neural responses. Hence, it is essential to control these potentially confounding variables (Anderson et al., 2006; Hart et al., 2012).

The current study introduces a naturalistic audio stimulus database that meets the specific requirements of cognitive neuroscience for short, recognizable, and emotionally neutral stimuli. Further, hierarchical categorization of these stimuli is clarified with a novel data-driven approach. To achieve this, 291 audio stimuli were collected from a wide range of sources. These audio clips were rated by participants on emotionality, arousal, and recognizability, and subsequently freely described by typing what they believed the sound to be. These responses were embedded and clustered using an unsupervised machine-learning algorithm to derive a participant-grounded organization of auditory object categories. This database is designed for cognitive neuroscience and neuroimaging research, as well as in the broader field of behavioral sciences.

## 2 Methods

### 2.1 Stimuli Curation

Stimuli were obtained from two repositories: BBC Sound Effects (BBC, 2018) and Freesound.org. The BBC Sound Effects library provides sounds licensed for non-commercial use, whereas Freesound.org distributes audio under various Creative Commons licenses, including CC0 (unrestricted use), Attribution (credit required), and Attribution–Noncommercial (credit required, non-commercial use only). A complete list of credited Freesound.org (Freesound, 2005) contributors is provided in the supplementary materials.

The sounds were selected to adhere to predefined criteria: each sound had to have a duration of less than 1.5 s, or be amendable to trimming of 1.5 s without loss of its characteristics. Sounds that were artificial, contained high levels of static noise (e.g., audible background white noise), were discontinuous (e.g., footsteps), or were alarming (e.g., an emergency siren) were excluded from the final database. The final stimuli set contained 291 sounds.

To facilitate balanced stimulus presentation in subsequent validation tasks, the 291 sounds were initially divided into 12 provisional semantic categories, each containing 21 - 30 recordings. These labels were organizational placeholders intended solely to ensure diversity of auditory sources during the experiment. Importantly, they did not reflect an assumed semantic or emotional structure. Instead, the final categorical organization of the stimulus set was derived empirically from participant responses collected during the validation experiment.

### 2.2 Stimuli Processing

All sounds were edited to a uniform 1.5-second duration, with fade-in and fade-out effects applied to minimize variability in onset. The audio files were then converted to a mono (i.e., centered in perceived location, to eliminate spatial localization cues), compressed to ensure the loudness of the audios stays consistent over the 1.5 seconds (i.e., bringing the quietest and loudest parts of the audio closer to each other), and normalized to −10 dB relative to full scale (i.e., bringing the average perceived loudness of the audio to −10 dB relative to full scale). Finally, any sounds exhibiting clipping or unnatural alterations were excluded.

### 2.3 Stimuli Norming Experiment

#### 2.3.1 Participants

A total of 445 participants were recruited via Prolific. Eighty-four did not complete the task or failed attention checks, resulting in a final sample of 361 participants (mean age = 29.67 ± 5.96; 113 females, 237 males, 11 others). Eligibility criteria included: age between 18 and 40 years, native or fluent English proficiency, normal hearing, no color blindness, and no self-reported mental or neurological problems. Only individuals using headphones were permitted to participate.

Participants were randomly assigned to one of three groups, each rating approximately one-third of the database. All participants provided electronic informed consent. The study was approved by the Ethics Committee of the University of Muenster (2024-40-AZ) and conducted in accordance with the Declaration of Helsinki. Compensation was £10.5/hour.

#### 2.3.2 Behavioral Standardization Task

The task was implemented using Psychtoolbox (Brainard, 1997; Pelli, 1997). Audio clips were presented consecutively, and participants were instructed to rate three dimensions of each clip: arousal, emotionality, and recognizability. For each rating, participants used a mouse-operated continuous slider bar with pictorial anchors at either end to illustrate the scale extremes (e.g., a confused face with a question mark for the low end of the recognizability scale). The format was adapted from the Self-Assessment Manikin (SAM), selected for its established reliability and low cognitive demand (Bradley & Lang, 1994).

At the beginning of the task, participants were provided with the following instructions. For arousal: “Please rate how much the audio startled you by moving the mouse to the left if the sound was not startling or to the right if the sound was startling. The amount of change in the position of the toggle on the screen should reflect your degree of confidence in the response.” For emotionality: “Please rate how the audio made you feel by moving the mouse to the left if you felt negative after hearing it (e.g. disgusted, sad, or angry) and to the right if you felt positive after you heard the sound (e.g. curious or happy). The bigger movement corresponds to stronger emotions.” For recognizability: “Assess how easily recognizable the sound was to you by moving the mouse to the left if the sound was unrecognizable and to the right if the sound was easily recognizable (e.g., keys jingle). The amount of change in the position of the toggle should reflect the degree of your confidence in the response.”

Following the ratings, participants were instructed to type one or two words describing the presented sound. The on-screen instructions read: “Please use the keyboard to type one word or two to describe what you think the audio was. Please try to type this as quickly as possible and limit your input to one or two words at most. Please try to keep your guess as general as possible. For example, if the sound is of tree leaves rustling, please write plants and not tree leaves.” This open-ended format was chosen to capture naturally occurring semantic categories rather than constrain responses to predefined options.

The experimental sequence began with a fixation cross, which was displayed for 600 ms, jittered by 50 ms, followed by a sound (1,500 ms), then a second fixation cross (600 ms, jittered by 50 ms). After the second fixation cross, the three rating scales were shown in a fixed order (arousal, emotionality, and recognizability). For each scale, participants had 6 seconds to respond. Finally, the open-ended question was presented, with a maximum of 8 s to respond. Sounds were presented in a pseudorandomized order such that each author-defined category was represented at least once every 12 trials.

Interleaved between the main trials, 10 attention trials were presented throughout the experiment. The attention trials were pseudorandomized such that at least one occurred within each block of 10 sound trials. In the attention trials, participants were required to correctly select the color of a square presented in the middle of the screen from three options. Participants were required to answer correctly on at least 70% of attention trials; otherwise, their data were excluded. Participants had 60 s to answer the attention question; failure to respond within the time led to immediate termination of the experiment and exclusion from the study.

A rest period was included after 50 standard (sound) trials, corresponding to the midpoint of the experiment. Participants were instructed to take a 5-minute rest if they wished. During this rest phase, a timer appeared on the screen to inform participants of the remaining time before the experiment automatically resumed. Participants were able to terminate the rest by pressing the space bar at any time during this 5-minute period.

A practice block was administered at the start to familiarize participants with the task. At the beginning of the practice phase, an audio clip was played, and participants were asked to adjust the volume until it reached a comfortable level to avoid further adjustments during the experiment. Since all the sounds were standardized for loudness, a single volume adjustment at the beginning was sufficient. After that, the example audio was played again, and participants were asked to rate the audio on the three dimensions and type a one or two-word description. The example audio used during the practice phase was unrelated to any experimental stimuli. After this practice trial, the main task began.

## 3 Data Records

Data can be found on the Open Science Framework website. The data is organised in five different folders as follows:

- Meta_Data: This folder contains the Meta_Data.xlsx worksheet, which lists the source of each audio file and its description from the original audio source.
- Processing_results: This folder contains the Audio_Categorization.xlsx and Ratings_Norm.xlsx worksheets.
  - Audio_Categorization.xlsx: This file contains the results of the two-step k-means algorithm that was run to empirically categorize the sounds using the data given by the participants
  - Ratings_Norm.xlsx: This file contains the normalized average ratings of each of the participants for each of the sounds
  - Raw_Data: This folder contains the Raw_Data_Ratings.xlsx and Raw_Data_Text.xlsx worksheets, which contain ratings and participants’ text guesses from the standardization study. Each participant was given an arbitrary number between 1 and 361 here.
- Sound-Standardization.zip: This folder contains a documented version of the analysis script, preprocessed data, and analysis results. An overview of the code is available in the README.md file.
- Sounds: this folder contains the main audio files, divided into 12 subfolders, each with 23 to 29 audio files. It is important to note that the authors provided the folder names assigned to these folders for ease of indexing and file handling, and to ensure a sufficient variety of sounds was presented to participants during the standardization process. The empirically derived sound categories are in the Audio_Categorization.xlsx file. The audio files are named with two letters indicating the folder they are in and a unique identifier. This letter-number combination will be used in all other .xlsx files, as well as the analysis scripts found in the Code Availability section.

## 4 Technical Validation

### 4.1 Rating Data Quality

Participants who completed fewer than 80% of rating scales were excluded from rating analyses (33 exclusions). For each sound, mean arousal, emotionality, and recognizability ratings were computed and z-normalized.

Normality was assessed using the Shapiro–Wilk test with Lilliefors correction. Arousal ratings were normally distributed (mean = 5.00 ± 0.70, *D*(291) = 0.03, p = .50), whereas recognizability (mean = 6.75 ± 1.22, *D*(291) = 0.09, p < .001) and emotional valence (mean = 5.70 ± 0.95, *D*(291) = 0.06, p = .03) were not. Both non-normal scales exhibited positive skew: recognizability values indicated high participant confidence in identifying sounds, and emotional valence suggested slightly more positive affective responses overall. Figure 1 displays distributions of the three rating dimensions.

**Figure 1.**
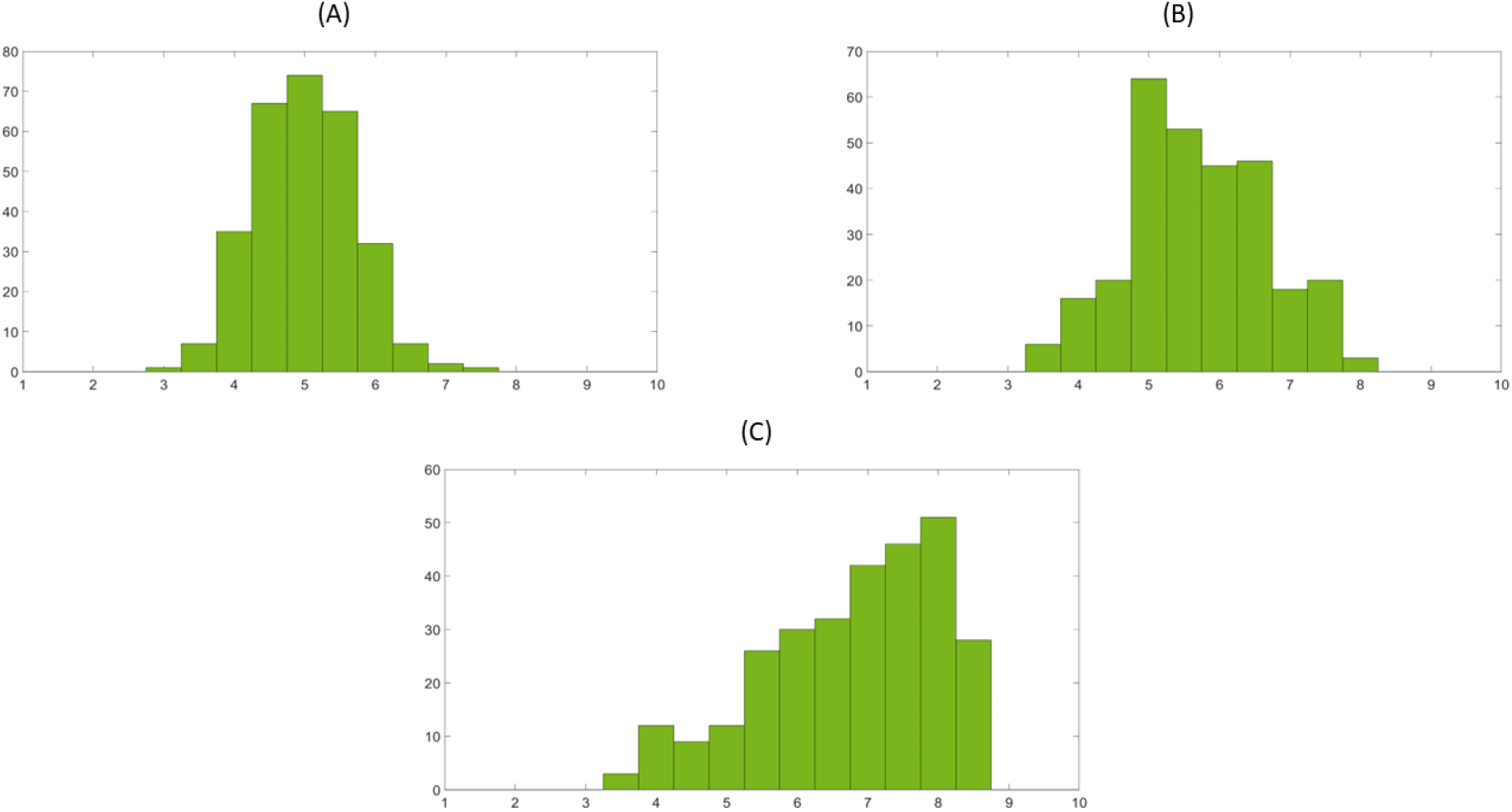
Histogram of the average rating of the sounds’ (A) arousal, (B) emotionality, and(C) recognizability.

To test the inter-rater reliability a two-way random, average score ICC, which measures inter-rater consistency was calculated. *ICC(2, k)* values were .86, .97, and .95 for arousal, recognizability, and emotionality respectively. These results suggest that the mean ratings for each sound provide a very stable and reliable measure of the intended psychological dimensions.

### 4.2 Text Processing Pipeline

Free-text responses were cleaned by removing non-sensical entries and stop-words using MATLAB’s built-in list, supplemented with: voice, noise, sound, noises, sounds, voices, and answer. Typos in the text responses were corrected using MATLAB’s native autocorrection tools (“correctSpelling” function). The MATLAB script including the correction procedure is included in the Code Availability section.

We used the participants’ guesses to categorize the sounds. To achieve this, we implemented a two-step procedure. In the first step, we applied semantic embedding (Pennington et al., 2014), a technique commonly employed in natural language processing (NLP) (Asudani et al., 2023; Peters et al., 2018). In this approach, each word is mapped to a multidimensional space to quantify its relationship with all other words in the language’s lexicon. To understand this step, one might consider mapping the words “queen” and “king” into a two-dimensional space. Although these two words are close on one dimension (related to “royalty”), they are different in another dimension (e.g., in “gender”). Using this quantification, one can then categorize the words. Notably, any lexicon will have more than two dimensions, and these dimensions are not as easily interpretable as in our example, since they lack a predefined semantic meaning.

The guesses in our study were transformed into a 200-dimensional GloVe semantic embedding space, trained on ~27 billion words from Twitter (Pennington et al., 2014). After cleaning the participant data, each answer was entered into the first-step analysis as a separate guess, regardless of the sound associated with it. Each of the 291 sounds received around 126 guesses in total. In the first step, we only clustered unique guesses to maintain a balanced category density and variance and thus preventing the formation of clusters that contain only very few sounds with very similar guesses across most participants. In essence, having the same guesses for some sounds would force the k-means algorithm to find categories with very little variance which would eventually lead the algorithm to categories each sound on its own.

After mapping the guesses into the embedding space, a k-means algorithm was used to categorize them. To determine the optimal number of clusters for the first-step k-means algorithm, the cluster number with the maximum silhouette score was selected. The silhouette score is a standard measure in unsupervised machine learning for selecting the optimal number of clusters and avoiding over- and underfitting (Dalmaijer et al., 2022; Layton et al., 2012; Pavlopoulos et al., 2025).

Afterward, for each sound, the number of responses assigned to each cluster was counted. Notably, we used the complete set of guesses rather than just the unique ones to classify sounds based on the full distribution of participant responses. Therefore, repeated guesses were retained to weight clustering outcomes, rather than reducing the analysis to unique guesses. A sound was classified into a cluster if the plurality of responses for that sound fell into that cluster. The k-means algorithm is subject to stochasticity due to the noise in the data. To decrease the impact of noise on the final classification, we conducted clustering 1,000 times using different random seeds. For each run, the optimal number of clusters was identified using the maximum silhouette score and then applied in the k-means algorithm with the same random seed to categorize participants’ guesses.

Finally, we created a representation similarity matrix (RSM) for the sounds based on the clustering results of the guesses for each sound. To do so, a 291 × 291 RSM was constructed; each entry in this matrix represented the normalized number of times two sounds were categorized in the same cluster (i.e., if *aij =* 10/1000, it means that sounds *i* and *j* were bundled in the same cluster 10 of 1000 runs). This step closely resembles ensemble learning in supervised learning (Sharma & Borah, 2024).

In the second step, another k-means algorithm was applied to RSM to cluster the sounds. Again, silhouette scores determined the optimal number of clusters.

Notably, the second k-means clustering was applied only once, as the first-step output yielded a single RSM with an acceptable signal-to-noise ratio.

Finally, for visualization, an agglomerative hierarchical clustering algorithm was run on the combined RSM. The resulting dendrogram illustrates the relationships between individual sounds within categories and between categories overall.

### 4.3 Text Processing Results

For the first step of sound classification, silhouette scores were calculated to determine the optimal number of clusters for the first-step k-means algorithm. Silhouette scores were computed for 5 to 25 clusters. Figure 2 shows the histogram of the chosen optimal number of categories based on the maximum silhouette score in each run. The histogram shows a clear tendency for the algorithm to cluster the responses into 10 categories.

**Figure 2.**
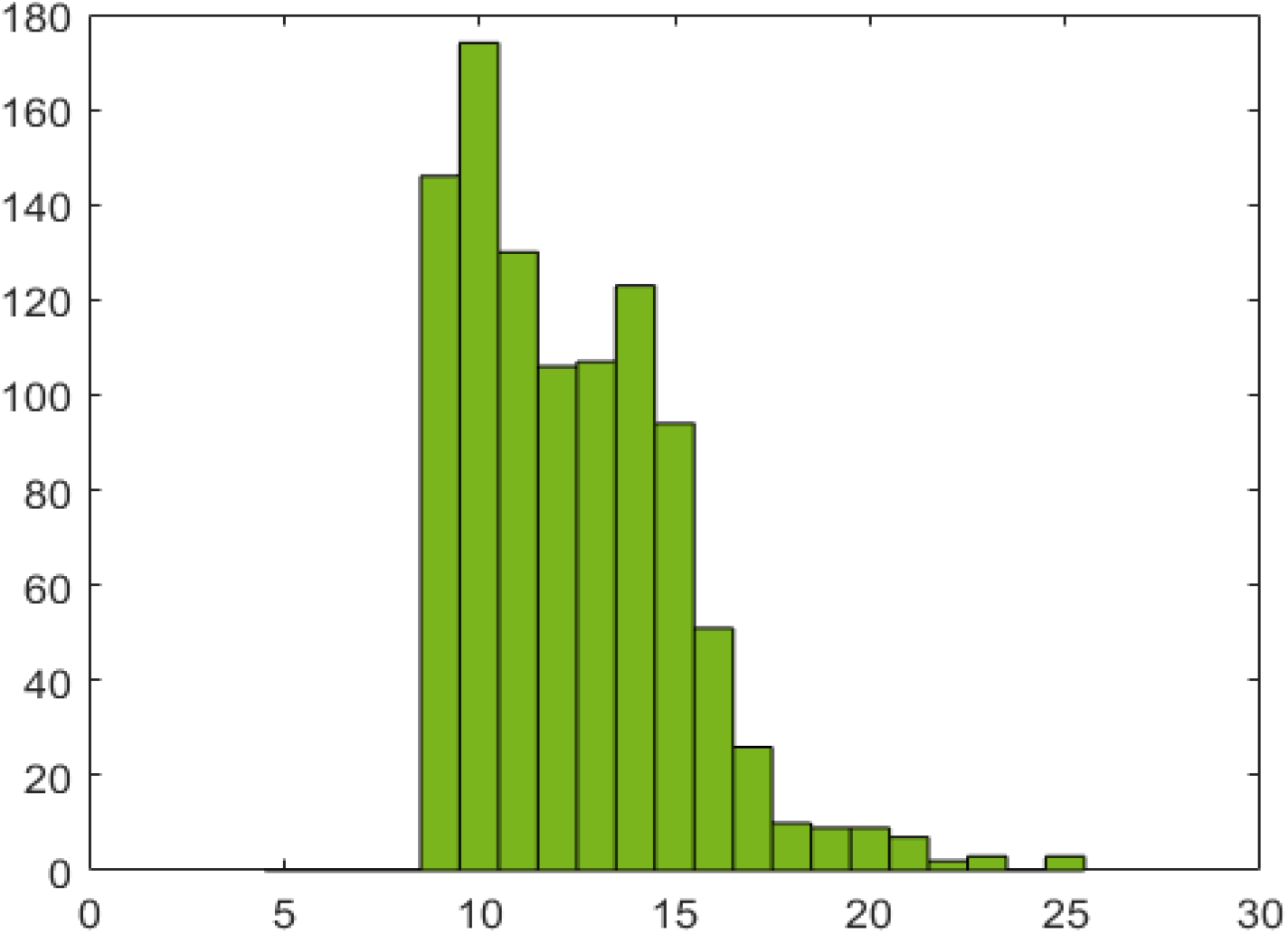
Histogram of the optimal number of categories based on the maximum silhouette score in the first step.

In the second step, the k-means algorithm classified sounds into 10 clusters. The 10 clusters were chosen based on the maximum silhouette score computed across cluster solutions ranging from 1 to 25 (Figure 3). The average silhouette score for the 10 clusters was 0.89, indicating high reliability of the k-means solution (Kaufman & Rousseeuw, 2005).

**Figure 3.**
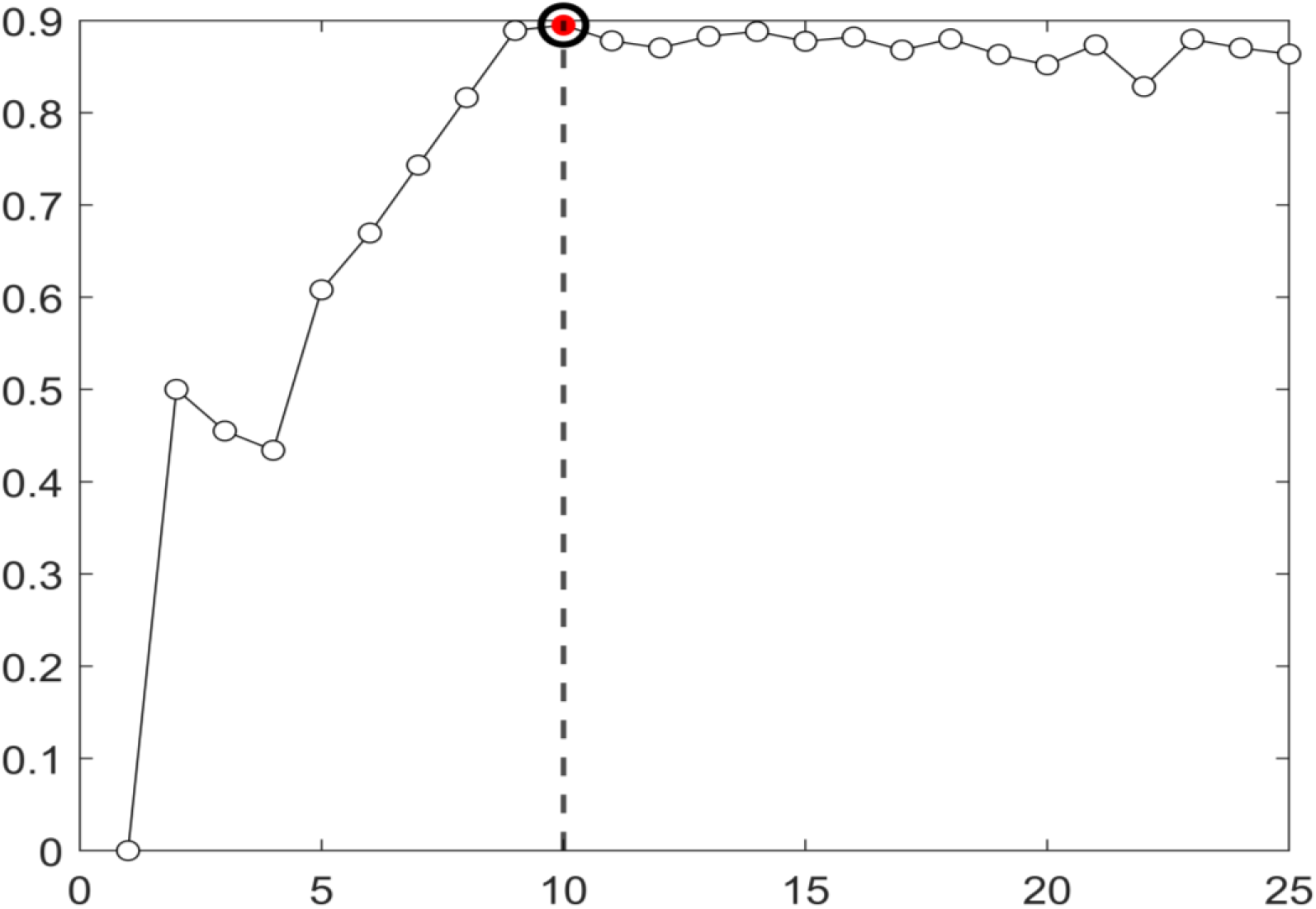
Average silhouette score for the second k-means step clustering sounds based on the combined RSM.

Table 1 summarizes the main results of the analysis, showing how each sound was classified into its respective category. The sound labels (e.g., ME01: Mechanical & Electronic sound 1) reflect the original grouping defined by the authors (see the Data Records section for a full list of the author-defined categories). Each column in table instead represents how the sounds were classified based on the participants’ responses. As shown, most sounds clustered as anticipated by the authors, albeit with a few exceptions, such as the speech category merging with bodily sounds (I.e., SP and BO categories).

**Table 1.** Categorizations of Sounds.

| Category 1 | Category 2 | Category 3 | Category 4 | Category 5 | Category 6 | Category 7 | Category 8 | Category 9 | Category 10 |
| --- | --- | --- | --- | --- | --- | --- | --- | --- | --- |
| BI10 BI11 | CG43 CG44 | BO01 BO02 BO03 | BS21 BS23 BS24 | BS26 MD01 | NW13 | BI12 BI14 BI15 | BS01 BS02 | CG01 | BO30 CG16 CG17 |
| BI16 BI13 | CG45 ME10 | BO04 BO05 BO07 | ME14 ME19 ME25 | MD12 MD13 | NW15 | BI18 BI02 BI20 | BS03 BS04 | CG02 | CG31 CG32 CG36 |
| BI38 MA01 | SO01 SO05 | BO08 BO09 BO10 | ME34 ME36 NW16 | MD14 MD17 | NW18 | BI24 BI25 BI27 | BS05 BS06 | CG14 | CG37 CG38 CG39 |
| MA03 MA06 | SO07 SO08 | BO11 BO15 BO16 | OA22 SO26 TR01 | MD18 MD02 | NW26 | BI29 BI03 | BS07 BS10 | CG33 | CG40 CG41 |
| MA07 MA08 | SO09 SO10 | BO17 BO18 | TR02 TR03 | MD20 MD21 | NW27 | BI30 BI32 | BS11 BS12 | CG35 | CG42 CG46 |
| MA09 MA11 | SO11 SO12 | BO19 BO22 | TR04 TR05 | MD22 MD23 | NW30 | BI33 BI34 | BS13 BS14 | NW01 | CG48 CG49 |
| MA12 MA14 | SO13 SO16 | BO24 BO25 | TR06 TR07 | MD28 MD30 | NW31 | BI36 BI39 | BS15 BS16 | NW02 | CG50 CG51 |
| MA15 MA16 | SO18 | BO26 BO27 | TR11 TR13 | MD31 MD32 | NW32 | BI05 BI08 | BS17 | NW03 | ME03 ME04 |
| MA17 MA18 | SO21 | BO28 BO29 | TR14 TR15 | MD33 MD35 | NW34 | OA01 OA02 | BS18 | NW04 | ME05 ME06 |
| MA19 MA20 | SO22 | SO24 SP03 | TR16 TR17 | MD37 | NW35 | OA03 OA04 | BS19 | NW05 | ME07 ME08 |
| MA21 MA23 | SO23 | SP05 SP07 | TR18 TR19 | MD39 | NW40 | OA05 OA06 | BS20 | NW06 | ME09 ME12 |
| MA24 MA26 | SO25 | SP08 SP11 | TR20 TR24 | MD04 |  | OA08 OA09 | BS22 | NW07 | ME20 ME26 |
| MA27 MA28 | SO27 | SP13 SP14 | TR25 TR26 | MD40 |  | OA11 OA12 | BS25 | NW08 | ME32 ME33 |
| MA29 MA30 | SO29 | SP15 SP16 | TR27 TR28 | MD05 |  | OA13 OA14 | ME01 | NW09 | ME35 |
| MA31 OA24 | SP01 | SP20 SP21 | TR29 TR30 | MD08 |  | OA15 OA16 | ME02 | NW10 | NW36 |
| OA25 OA26 | SP02 | SP23 SP31 | TR31 TR32 | MD09 |  | OA17 OA18 | ME21 | SO17 | NW39 |
| SP26 | SP06 | SP34 | TR33 TR34 | SP19 |  | OA19 OA20 | ME22 | SO28 | SO04 |
| SP33 | SP22 | SP35 | TR36 TR37 | SP32 |  | OA21 OA23 | ME23 | SO30 | SO06 |

*Note.* Name of the sounds represent the original author-defined categories as follows: ME: Mechanical & Electronic; CG: Household Items; BO: Bodily Sounds; NW: Natural & Water Sounds; BS: Bells & Sirens; MD: Music & Drums; SO: Sports Sounds; SP: Speech Sounds; TR: Transportations; MA: Mammals; BI: Birds; OA: Other Animals

Furthermore, Figure 4 displays a scatter plot of the sounds (i.e., the combined similarity matrix) and their categorization from the second step, after reducing the matrix to two dimensions using the t-SNE algorithm. This is a non-linear dimensionality reduction algorithm that prioritizes preserving local neighborhoods over global geometry, allowing clearer visualization of clusters (van der Maaten et al., 2008).

**Figure 4.**
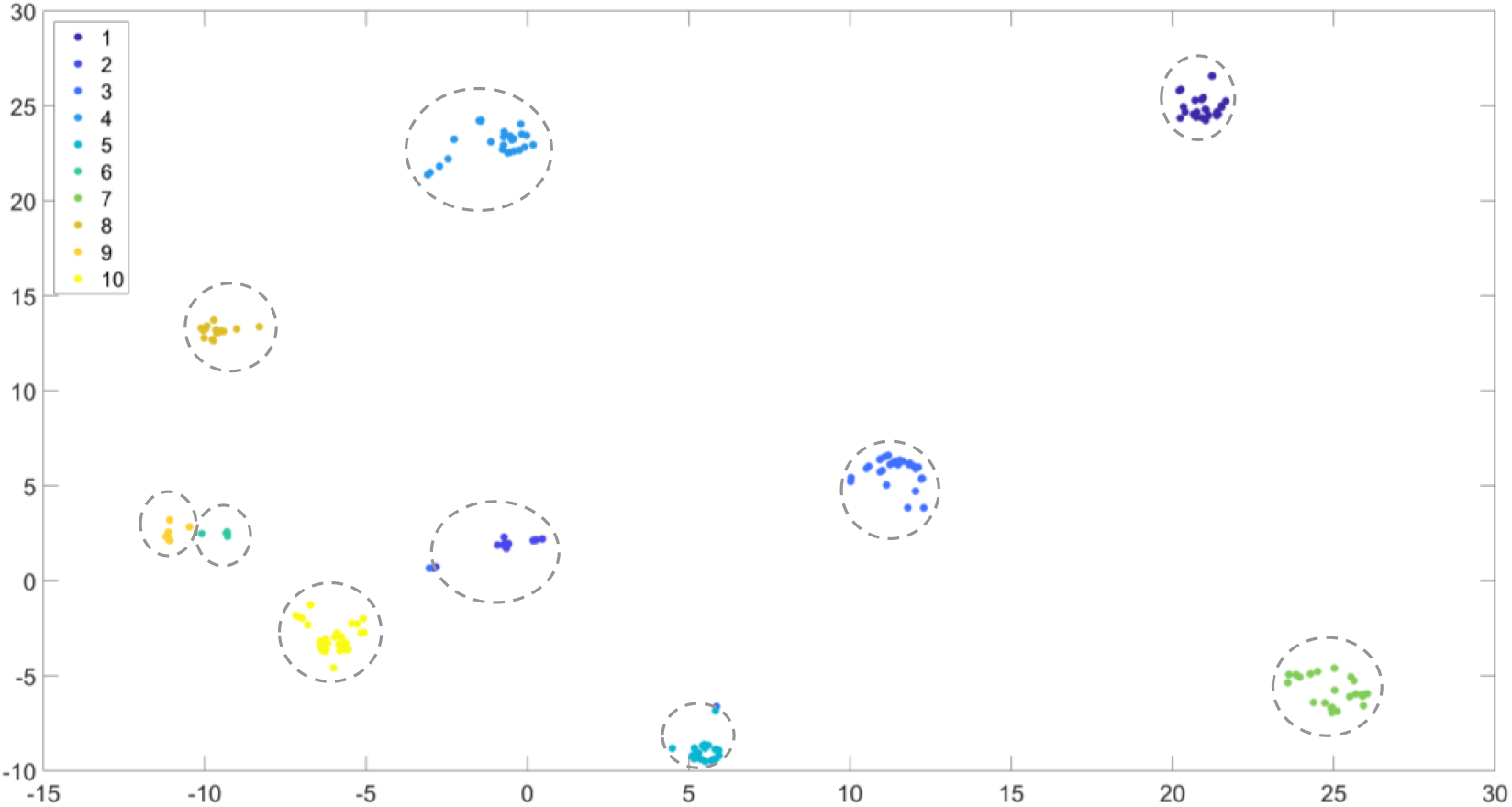
Scatter plot of the sounds based on t-SNE projections. Numbers in the legend correspond to the category numbers in Figure 3.

Lastly, to understand the full relationship between sounds and their categories, Figure 5 shows a complete dendrogram from hierarchical clustering applied to the combined RSM.

**Figure 5.**
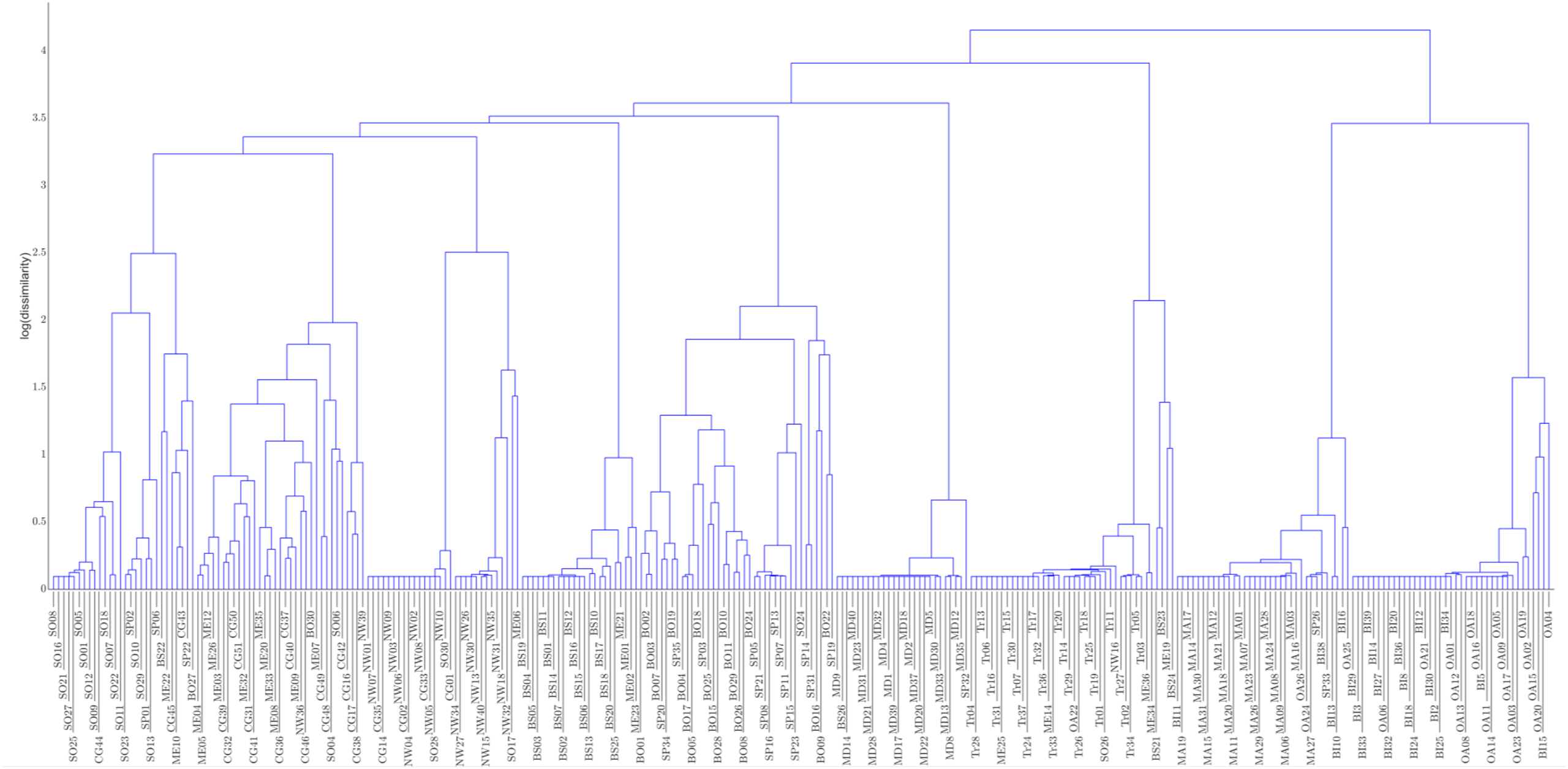
Full Dendrogram of the Sounds.

## 5 Usage Notes

Researchers can use the audio subset based on their research question. ‘Audios_Rating.xlsx’ contains the average ratings of the participants for each of the audio. Audio_Categorization.xlsx contains the labels assigned to each audio. Alternatively, researchers could use the raw rating and text data provided to run their own analyses or train their own machine learning models.

## 6 Data and Code Availability Statement

The data and code used in this study are available at the following link: https://osf.io/7wqnm/overview?view_only=af0f2b4608c6407497daf8360d9a3710

## Funding

No funding was obtained for this study.

## Notes

### Competing Interest Statement

The authors have declared no competing interest.

